# Hypoarousal non-stationary ADHD biomarker based on echo-state networks

**DOI:** 10.1101/271858

**Authors:** D. Ibanez-Soria, A. Soria-Frisch, J. Garcia-Ojalvo, Jacobo Picardo, Gloria García-Banda, Mateu Servera, Giulio Ruffini

## Abstract

Attention-Deficit Hyperactivity Disorder (ADHD) is a neurodevelopmental disorder characterized by inattention, hyperactivity and impulsivity. It is one of the most commonly diagnosed neurodevelopmental and psychiatric disorders of childhood and therefore presents a very high prevalence rate. However the high rate of ADHD misdiagnosis makes the discovery of neurophysiological ADHD biomarkers an important clinical challenge. This study proposes a novel non-stationary ADHD biomarker based on Echo State Networks to quantify EEG dynamical changes between low attention/arousal states (resting with eyes closed, or EC) and normal attention/arousal states (resting with eyes open, or EO). Traditionally, EEG biomarkers have revealed an increase in stationary power in the theta band along with a decrease in beta, with these frequencies largely accepted to be altered in the ADHD population. We successfully verify the hypothesis that measured differences between these two conditions are altered in the ADHD population. Statistically significant differences between a group of ADHD subjects and an aged-matched control population were obtained in theta and beta rhythms. Our network discriminates between EO/EC EEG regimes in the ADHDs better than in controls, suggesting that differences in EEG patterns between low and normal arousal/attention states are larger in the ADHD population.

## 1. INTRODUCTION

Attention deficit hyperactivity disorder (ADHD) [1] is a chronic, pervasive childhood disorder characterized by low frustration tolerance, excessive impulsivity, distractibility, and inability to sustain attention and concentration [2,3,4]. It is one of the most commonly diagnosed [2] and investigated [4] neurodevelopmental disorders of childhood. Three ADHD presentations or subtypes can be diagnosed from 9 symptoms of inattention (IN) and 9 symptoms of hyperactivity/impulsivity (HI) using DSM-IV [5]: Combined (6 or more IN and 6 or more HI symptoms), predominantly inattentive (6 or more IN and less than 6 HI symptoms), and predominantly hyperactive/impulsive (6 or more HI and less than 6 IN symptoms). ADHD has a high prevalence rate that is estimated between 5% and 7% [6, 7].

Although ADHD is currently considered a neurodevelopmental disorder [8], the diagnostic criteria continue to be based primarily on subjective behavioral measures derived from parent and teacher reports, interviews, or direct observation. Therefore, as Crippa et al. (2017) point out, the diagnosis is heavily based on the experience and practical knowledge of clinicians. This has at least two important consequences. First, there is a social and scientific concern about the reliability and variability of this approach to diagnosis, and the potential for high probability of a misdiagnosis [9,10]. And, second, although some authors consider that brain structural and functional deficits have been proven to be associated with ADHD [10], there is not a clear consensus about this point. Therefore, finding solid evidence of neuropsychophysiological dysfunction in ADHD has become one of the most relevant challenges in mental health research.

In recent years, electroencephalographic (EEG) measures in resting-state conditions have been widely used to monitor neurophysiological abnormalities in the ADHD population [11]. Most reported findings have shown that the ADHD population presents an increased power in fronto-central regions in low frequencies (typically in the theta band) [12, 13] with decreased power in fast frequencies (typically in the beta band) [14, 15]. Theta/beta ratio (TBR) has long been used as an ADHD biomarker [16]. The US Food and Drug Administration (FDA) approved the Neuropsychiatric EEG-Based ADHD Assessment Aid (NEBA®), which uses the theta/beta ratio of the EEG measured in the central EEG electrode Cz combined with a clinician’s evaluation to support the diagnosis of ADHD. NEBA cutoffs for analysis were pre-established and are different for adolescents and children [17,18]. However, not all recent studies could validate the usage of TBR, as a biomarker for diagnosing ADHD. Recent studies documented an insufficient accuracy for TBR and theta power in distinguishing children with ADHD from a control group [11, 19, 20]. Therefore the discovery of novel robust ADHD biomarkers remains a hot research topic.

EEG band-power assessments assume a large degree of temporal stability in brain oscillations. Typically, in EEG analysis the signal is split into short-time epochs that are considered to be pseudo-stationary. Band power is estimated at each epoch and subsequently averaged across them [21]. However, it is well known that the brain is a complex system that generates non-stationary EEG patterns of high dimensionality [22, 23]. Such dynamic, chaotic behavior advocates for the use of non-stationary EEG analysis techniques for EEG feature extraction and classification. Here we apply this approach to reexamine the hypothesis that ADHD is associated with a hypoaroused brain state, suggested by scientific evidence over the past decades. This hypothesis is based on the fact that arousal and attention are related and overlapping concepts [24]. Arousal acts as a modulator of attention levels, with changes in arousal followed by changes in attention [25]. Recent theories, such as the cognitive-energetic model [26], include the concepts of arousal, activation, and alertness as basic mechanisms in ADHD [27, 28, 29].

Following the hypoarousal ADHD theory, our hypothesis is that the magnitude of EEG differences between low attention/arousal states (during EC) with respect to normal attention/arousal states (during EO), may be altered in the ADHD population. We test here this hypothesis through the study of band-power stationary features, but also dynamical alterations in the temporal dynamics. To this effect, a novel approach based on recurrent neural networks (RNN) in its reservoir computing (RC) form (echo-state networks) is proposed here.

EEG Differences between eyes open (EO) and closed (EC) conditions have been largely reported in the alpha band. Arousal increase during resting state withEO with respect toEC has been associated with a global decrease of EEG alpha levels [30]. Alpha levels are in general substantially reduced in amplitude by eye opening [31] and its regime is characterized by a dominating oscillatory rhythm known as the peak alpha frequency (PAF). This rhythm is however not strictly monotonic, varying over a range of about 1 Hz [32]. Regarding EO-EC changes in other EEG bands, in resting-state EO conditions, reductions of absolute power levels in the delta, theta and beta bands between the two conditions have been reported in children. Power topographic changes across the scalp have been also observed in all bands [33].

We expect the EEG dynamic regimes, as measured through RC, during interleaving intervals of EC-EO, to differently discriminate in a different fashion for ADHD and healthy subjects. RC has been applied in the past to several EEG feature extraction and classification problems such as brain computer interfaces [34,35], epileptic seizure [36], prognosis in Parkinson’s disease [37] or event detection [38]. In a previous work [39] we have demonstrated RC capacities to well characterize complex dynamics among EEG channels when a dominating frequency in the EEG spectrum is elicited. Here we explore EO-EC discrimination capabilities within EEG frequency bands, that as alpha may be characterized by a dominant oscillatory regime. To the best knowledge of the authors, RC has never been applied in the field of ADHD biomarkers.

## 2 Reservoir Computing

Artificial Neural Networks (ANNs) are computational models inspired in the functioning of the brain [40, 41, 42]. Their structure consists of a network of interconnected artificial neurons also known as nodes or units. Artificial neurons transmit signals from one to another along the network simulating the biological synapse process. In practice, artificial neurons receive signals from connected neurons, with a fixed weight (*w*_*i*_) that is set during the network training process. The activation function of each neuron maps the sum of input weighted connections into the signal transmitted to other neurons. Typical activation functions in ANN are the rectified linear unit, the sigmoid, the hyperbolic tangent or the unit step. A representation of an artificial neuron is displayed in Figure 1A.

**Fig. 1.**
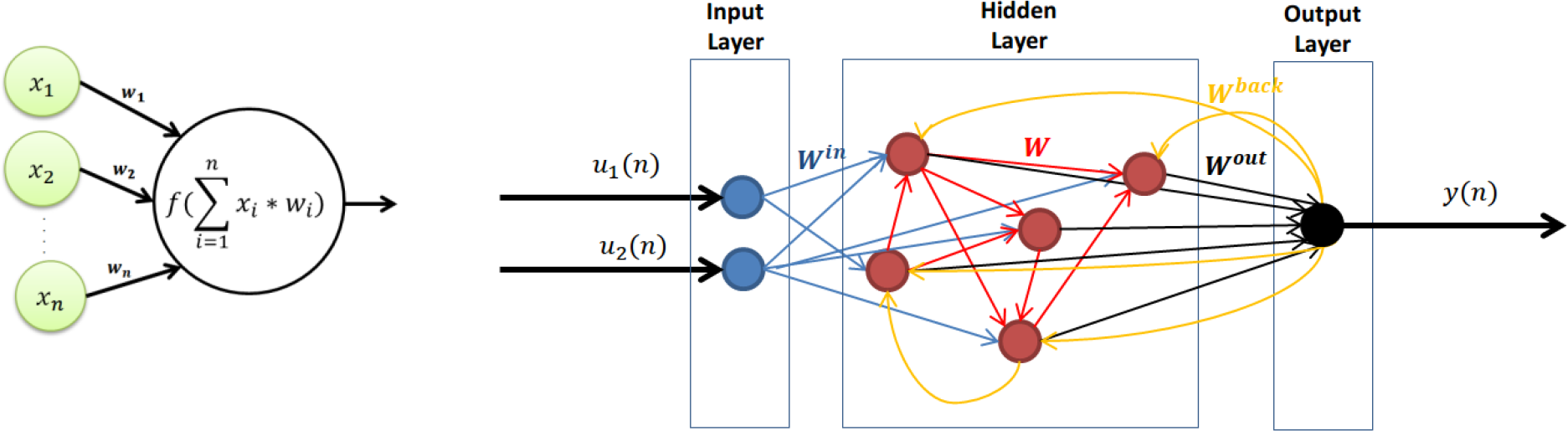
A) Artifical neuron and B) recurrent neural network Representation.

ANNs in general present three stratums of neurons: the input, hidden and output layer. Figure 1B represents a network similar to the one used in this work with 2 input units and one output unit. In the network two input signals *u*_*1*_(*n*) and *u*_*2*_(*n*) feed the two units of the input layer whose goal is to interface with input data. Input weights *W*^*in*^ map the input nodes into the hidden layer. Internal weights (*W*) interconnect hidden layer units while output weights (*W*^*out*^) map the hidden nodes into a single output node. Adding loops to the hidden layer transforms (*W*^*back*^) what would otherwise be a standard feedforward network (multilayer perceptron) into a recurrent network, and allows the model to encode time-resolved information and thus to incorporate memory, converting it into a dynamical system [43].

In the 2000s, reservoir computing (RC)–a new approach for training, understanding and using Recurrent Neural Networks– was proposed independently and simultaneously under the names of Echo State Networks (ESNs) [44] and Liquid State Machines [45]. Reservoir computing is based on the principle that certain structural properties of the network make the supervised training of all weights unnecessary. In particular, if the network obeys an algebraic property known as the echo state property (ESP) only readout connections need to be adapted in a supervised way. The untrained network, whose weights are fixed and randomly generated, is known as the dynamical reservoir (DR) and consists of input-scaling, back-projection and internal weights. The reservoir provides memory while nonlinearly expanding the input signal [46].

The echo-state property holds if the state of the network asymptotically depends only on the input signal, implying that initial conditions dependencies are lost progressively. In practice, input and back-propagation weights do not affect the echo state property, that only depends on internal weights of the hidden layer. In most applications if the spectral radius of internal weights, calculated as the largest absolute eigenvalue of the adjacency matrix of the reservoir, is kept below one, the ESP holds [46], although unlikely exceptions have been reported [47].

According to ESN best practices [46], the most important global-parameters that shall be optimized in order to achieve a good performance are the: 1) input scaling, 2) spectral radius, 3) leaking rate and 4) reservoir size. When scaling the network inputs, in practice the same input scaling factor is applied to *u*_*1*_(*n*) and *u*_*2*_(*n*). The input scaling drives the degree of non-linearity in the reservoir. Linear tasks require small input scaling factors while complex tasks demand larger input scaling values, easily saturating the nodes and thus transforming them into binary switches. The spectral radius governs the time scale of the reservoir, and thus determines how the influence of inputs remains in the system [48], and the leak rate determines the speed of the reservoir to update dynamics. The reservoir size is given by the number of internal units and is in general larger in ESNs compared to other neural network approaches [49]. It has to be large enough to learn the dynamics of the input signals, but not too large so it generalizes well with non-training data. The best approach for output weight training is ridge regression regularization, which removes the requirement of injecting noise in the network inputs to ensure a good generalization [46, 50].

Reservoir computing greatly simplifies the training of recurrent neural networks. The dynamical reservoir is randomly constructed according to selected hyperparameters. Once the DR weights are fixed, readout connections are learnt using a training input and a teacher-forced output. The network is thus able to efficiently perform tasks with complex temporal information with a low-training cost, since only the readout weights need to be trained.

## 3. METHODS

### 3.1 Participants

52 children aged 7-11 participated in this study. All subjects brought signed parental informed consent and were assigned to one of two groups: clinical diagnosed ADHD group or healthy controls. Children diagnosed with ADHD were recruited from clinical units specialized in pediatric disorders in Palma de Mallorca, Spain. To be considered for the ADHD group, children had to fulfill the following inclusion criteria: (1) being clinically diagnosed with ADHD by a specialist based on DSM-IV criteria (2) not having comorbidity problems of mental retardation, autism, bipolar or psychotic disorders, history of epileptic seizures or any other relevant medical disorder. 28 children were first recruited and assessed into this group, but during the data analysis 7 of them had to be rejected: 3 because they had taken medication 24h prior to the EEG assessment, and other 4 due to noisy EEG recordings. The final ADHD group thus included 21 children: 12 with combined ADHD subtype (11 males and 1 female) and 9 with inattentive subtype (3 males and 6 females).

Healthy controls were selected from standard school age-matched classrooms. The research team met with the schools’ principals and tutors, and gave them a dossier (including the informed consent) explaining the project. This dossier was sent home with the kids for their parents. Inclusion criteria for this group were: (1) not having any psychopathology diagnosis, neither mental retardation or learning disorders, (2) not showing behavioral problems nor learning difficulties in class (as asserted by their tutors) (3) not having major family problems that could interfere with their participation in the study. In the original control sample there were 43 children, but 13 had to be rejected at the final analysis: 7 of them due to academic problems, other 5 due to noisy EEG recordings, and the remaining one due to visual difficulties that prevented him from carrying out one of the experimental tasks. Thus, the final group of healthy controls consisted of a sample of 30 participants. The demographic characteristics of the experimental groups are summarized in Table 1 below.

**Table 1:** Experimental population description.

|  | ADHD (n = 21) | Control (n = 30) |
| --- | --- | --- |
| Age in years | M 9.6 (SD 1.4) | M 9.3 (SD 1.5) |
| Male/Female | 14 / 7 | 13 / 17 |
| ADHD C/I | 12 / 9 | - |
| ADHD Medication (Yes/No) | 6 / 15 | - |

### 3.2 Experimental Procedure

The most relevant findings in ADHD biomarkers have been obtained in fronto-central regions [51]. Following this, in this study we measured the brain activity of the participants in C3, Cz, C4, F3, Fz and F4 using six Ag/AgCl electrodes according to the 10/10 EEG standard positioning system [52]. EEG data were obtained with a Neuroelectrics Enobio^®^ recording system at a sampling rate of 500 Hz. The CMS/DRL electrical reference was placed in the right mastoid. The experimental protocol consisted of a 3-minutes resting-state eyes-open (EO) recording followed by a 3-minute eyes-closed (EC) recording. Participants were instructed to stay still looking at a fixation cross displayed in a computer screen at one meter-distance.

### 3.3 ESN-Based Dynamical Synchronization Metric

We now introduce our ESN-based approach aiming at quantifying within-subject dynamical differences between resting EO and EC states at specific frequency bands. The final goal is to employ such performance measure as a surrogate for the differences of EEG signal dynamics, and therefore as a biomarker for characterizing ADHD patients. The following paragraphs describe the signal processing and analysis pipeline that lead to the computation of the aforementioned biomarker.

Recordings are first filtered using a finite impulse response filter (FIR) at the following bands: theta1 (4-6 Hz), theta2 (6 −10 Hz), alpha1 (8 - 11 Hz), alpha2 (10 - 13 Hz), beta1 (13-20 Hz), beta2 (20-30 Hz), gamma1 (25-35 Hz) and gamma2 (35-45 Hz). The reference method has a substantial impact on potential measurements. Many reference strategies can be used in EEG analysis, including single electrode reference, linked-ears, linked-mastoids, ipsilateral-ear, contralateral-ear, bipolar references, the tip of the nose or weighted average electrodes subset among others [53]. Each modality has its own advantages and disadvantages. In practice, the electrode reference is chosen based on the electrode montage and characteristics of the feature to be computed. In this work the average reference of central electrodes C3, Cz and C4 has been used as reference. Frontal electrodes were not selected for referencing as they are likely affected by ocular artifacts. Given we are explicitly taking EO intervals into account, the use of frontal signals in the referencing process could distort the EEG dynamics of other channels.

After referencing, channels C3, Cz, C4, F3, Fz and F4 are split into 10-second epochs with no overlap. Epochs containing samples larger than 75uVs at any channel after detrending are rejected, as they are considered to be contaminated by artifacts. Since we are only interested in signal dynamics, each 10-second epoch is individually standardized to mean zero and standard deviation one, in order to remove the amplitude information. EC and EO standardized epochs are then sequentially concatenated creating a continuous EO-EC series for each channel and frequency band. The envelope of this series is then computed using the Hilbert transform [54].

ESN networks are fed with the previously defined interleaving temporal dynamics of EO and EC series coming from two EEG channels filtered at the same band. A teacher-forced signal with EO samples to 1 and EC samples to −1 is used for training, through which the network learns to distinguish between the two regimes. In previous works we have demonstrated that among other dynamics, ESNs are capable of detecting complex synchronization between two temporal time-series such as generalized synchronization [55]. We thus expect ESN to be capable of detecting synchronization variations among channels between the EO and EC regimes. This is why we will define the resulting biomarker as Channel Dynamical Synchronization Metric (CDSM). We set the ESN node activation function to the hyperbolic tangent, the spectral radius to 0.8, the input-scaling factor to 0.1, the leak rate to 0.5, and the number of units in the hidden layer to 500. The network output is low-pass filtered at 5 Hz in order to remove high-frequency components.

As explained above, we train the network to discriminate between EO and EC regimes for every combination of pairs of electrodes at one frequency band. To quantify these changes we compute the mean squared error (MSE) between the teacher-forced signal and the trained ESN output. We call this measure the channel dynamical synchronization metric (CDSM). This feature quantifies how well the network learns to discriminate between the EO and EC conditions at a certain band for a pair of electrodes. The dynamical connectivity index (DCI) of an electrode for a certain band is defined as the average DSM of its combination with every other electrode. In order to reduce the random effect introduced by the dynamical reservoir construction, we calculate the average DSM over five independent instantiations of the ESN, and use these replicates for further statistical analysis.

### 3.4 Stationary Analysis

We performed a comparative evaluation of the newly proposed approach with the conventional stationary spectral analysis, in order to obtain a better understanding of the ESN results and to explore stationary changes in the EEG rhythms. The average stationary spectral response and power at frequency bands defined in 2.3 is computed for each subject at EO and EC conditions using as reference also the average of C3, Cz and C4. The ratio of the average band power per channel between EC and EO conditions was calculated to measure the stationary differences between these two conditions at subject level.

### 3.5 Statistical Analysis

The performed statistical analysis aim at measuring if the samples under comparison, belonging respectively to ADHD and control groups are independent and therefore coming from populations with different distribution. According to the Kolgomorov-Smirnoff test [56] we could not guarantee a Gaussian distribution of all samples under test. Therefore, we used a nonparametric two-sided Wilkoxon rank-sum test [57], which unlike the t-test does not assume normal distributions, to identify statistical differences between the ADHD and control groups. The null hypothesis was rejected only if the obtained probability value was less than 5%. Statistical significance between groups is here represented by * for probabilities below 5%, ** for probabilities below 1% and *** for probabilities below 0.1%.

## 4. RESULTS

### 4.1. Dynamical Synchronization Metric Analysis

CDSM feature has been computed for every possible combination of pair of electrodes and frequency bands. Electrodes used in this study are presented in Figure 2A. Figure 2B shows a connectivity representation of the statistically significant p-values (p<0.05) obtained when comparing ADHD and control populations. We observe that the synchronization metric proposed here discriminates well between populations, especially in theta and beta bands. Figure 2C and 2D depicts the ADHD (blue) and control (red) CDSM grand averages and standard error of the mean (SEM) in the theta1 and beta1 bands. Given that the CDSM quantifies the error between the ESN output and the EO-EC ground truth, we can conclude that the network learned better differences between the two conditions in the ADHD population. This advocates for the presence of larger differences in dynamical EEG regimes in the ADHD group than in the healthy controls.

**Figure 2:**
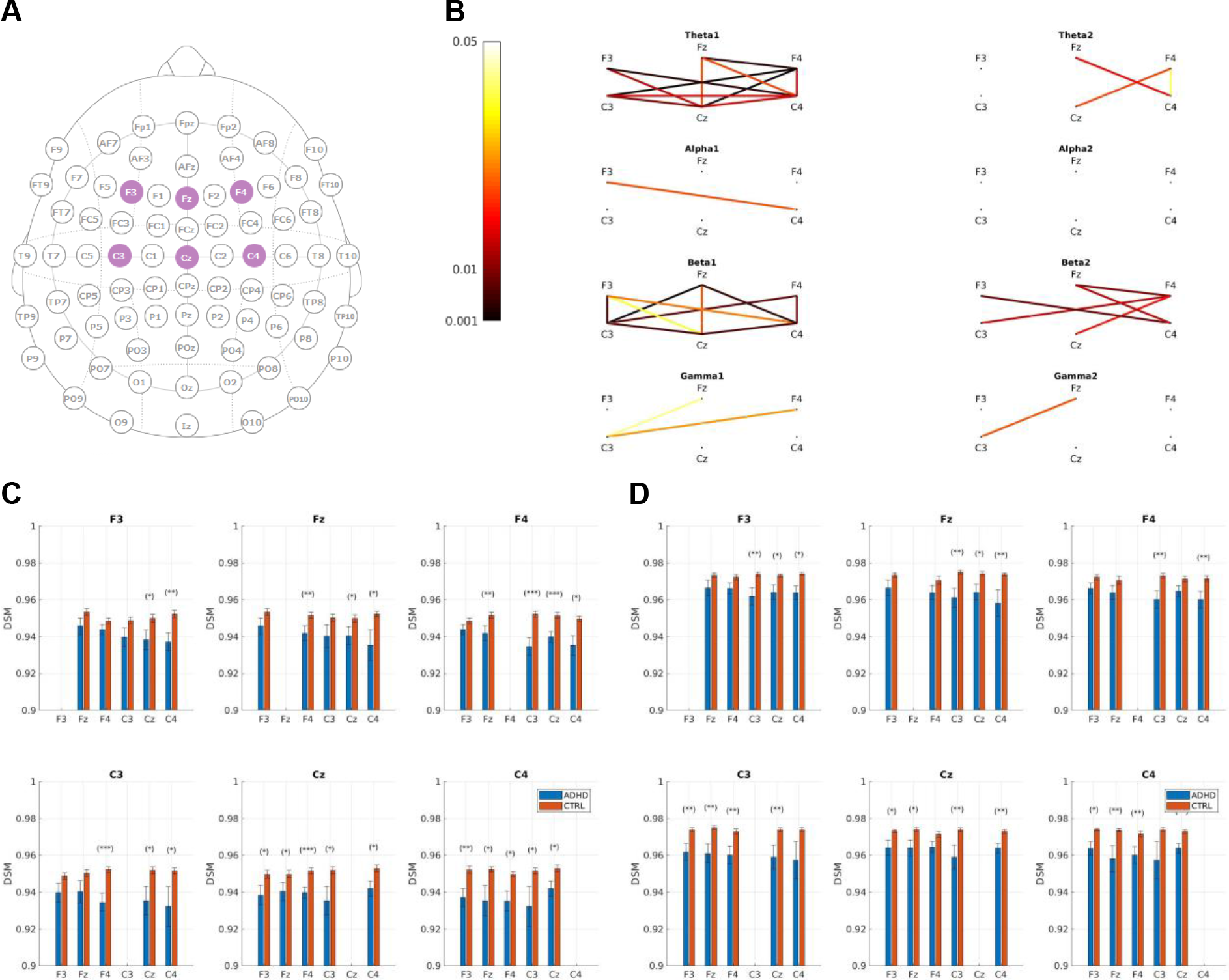
A) Position of the F3, Fz, F4, C3, Cz, C4 electrodes used in this study. 2) Connectivity CDSM representation of statistically significant p-values when comparing ADHD and control populations. 3) Theta1 CDSM grand averages and SEM. 4) Beta1 CDSM grand averages and SEM.

Figure 3 displays the grand average dynamical connectivity index together with the standard error of the mean. Results are consistent with previously discussed findings. The DCI is smaller in the ADHD population, showing a larger difference in its temporal dynamics between EC/EO conditions. Table 2A summarizes the statistical significance when comparing the two groups. As can be observed, differences between groups are representative in the theta1, beta1 and beta2 bands.

**Figure 3.**
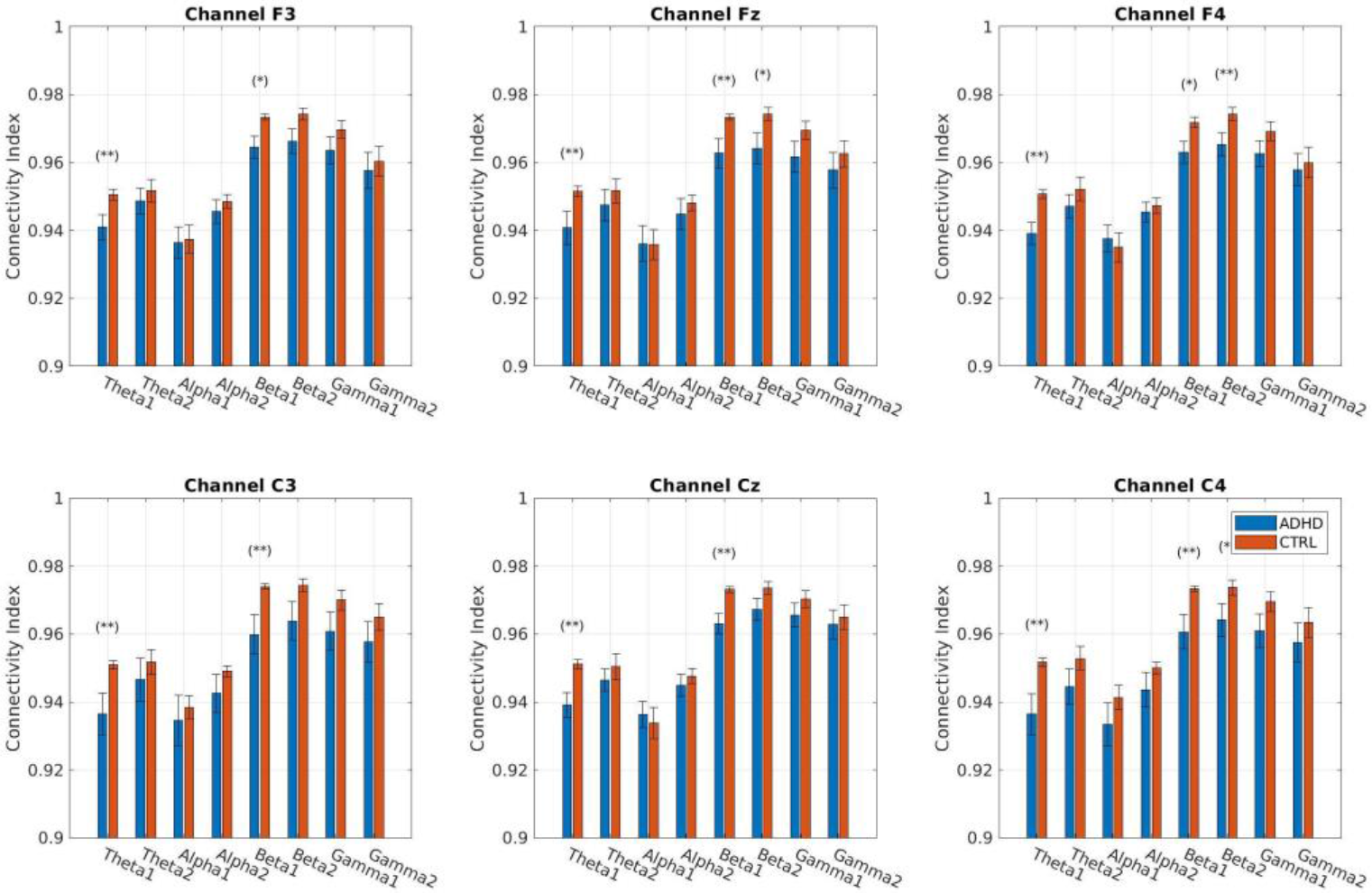
Dynamical Connectivity Index (DCI) grand averages and standard error of the mean in ADHD and control populations.

**Table 2.**
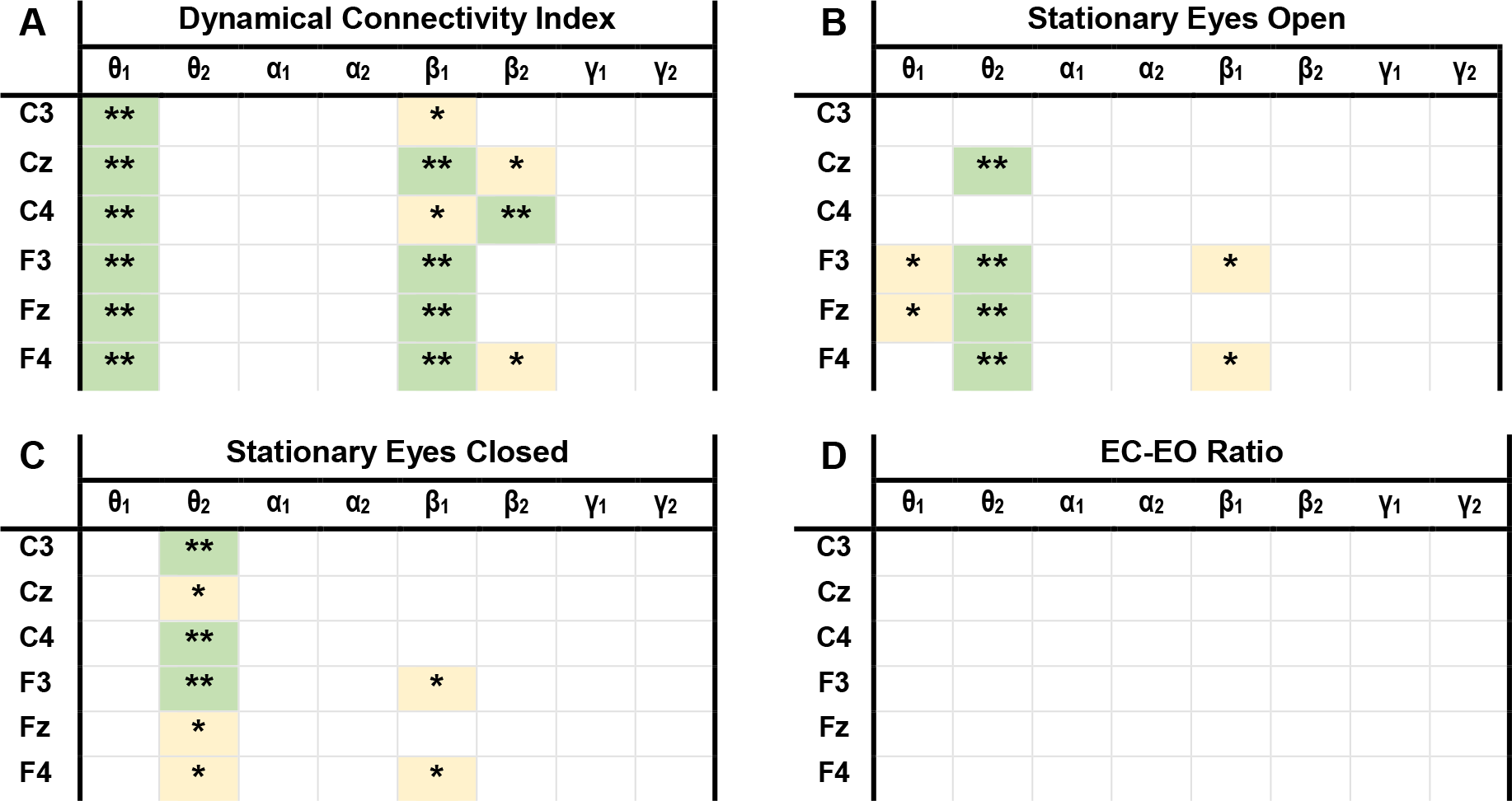
Statistical significance when comparing ADHD populations for A) Dynamical Connectivity Index, B) Eyes open stationary power, C) eyes closed stationary power, D) eyes closed-open stationary power ratio.

### 4.3. Stationary Band Power Analysis

Figure 4 represents the grand average spectral response together with the standard error of the mean in the 4-45Hz frequency interval. We can observe that the energy in theta and alpha increases in the EC condition for both the ADHD and control groups. When we look independently at each condition we observe an EEG slowing both in EO and EC. This effect is manifested a shift of the alpha peak frequency towards lower frequencies and an energy increase of low frequency components.

**Fig. 4.**
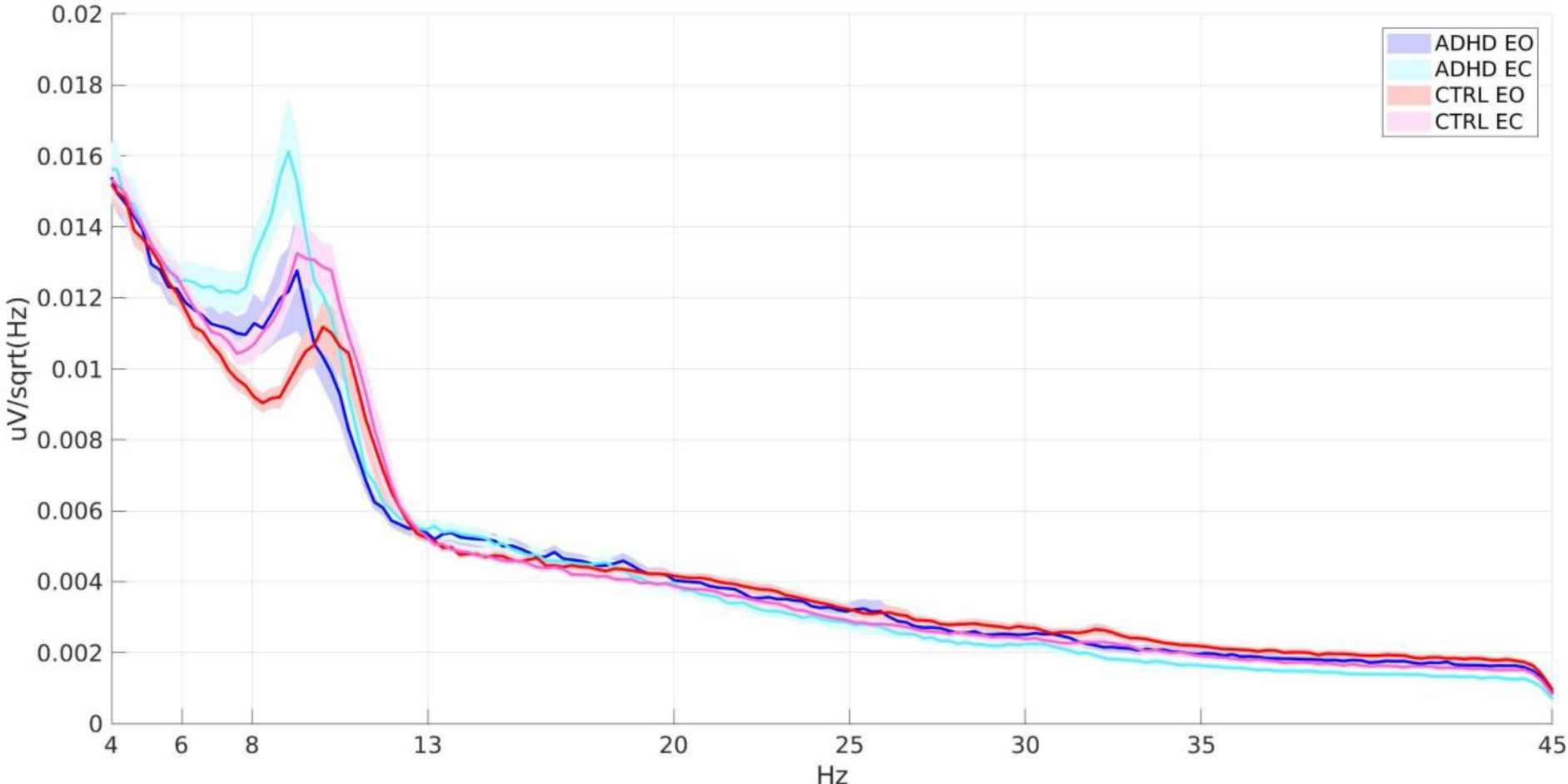
Grand average spectral response and standard error of the mean of ADHD EO, ADHD EC, Control EO and Control EC populations.

Figure 5A and B represent the power grand averages and standard error of the mean for the ADHD and control populations, for both the EO and EC conditions. We observe a power increase in low frequency bands theta and alpha in the ADHD populations for the two conditions. Statistically significant power differences between ADHD and Control groups are presented in Table 2B and 2C, where we can observe that theta2 delivers the most statistically significant differences both for EO and EC conditions.

**Fig. 5.**
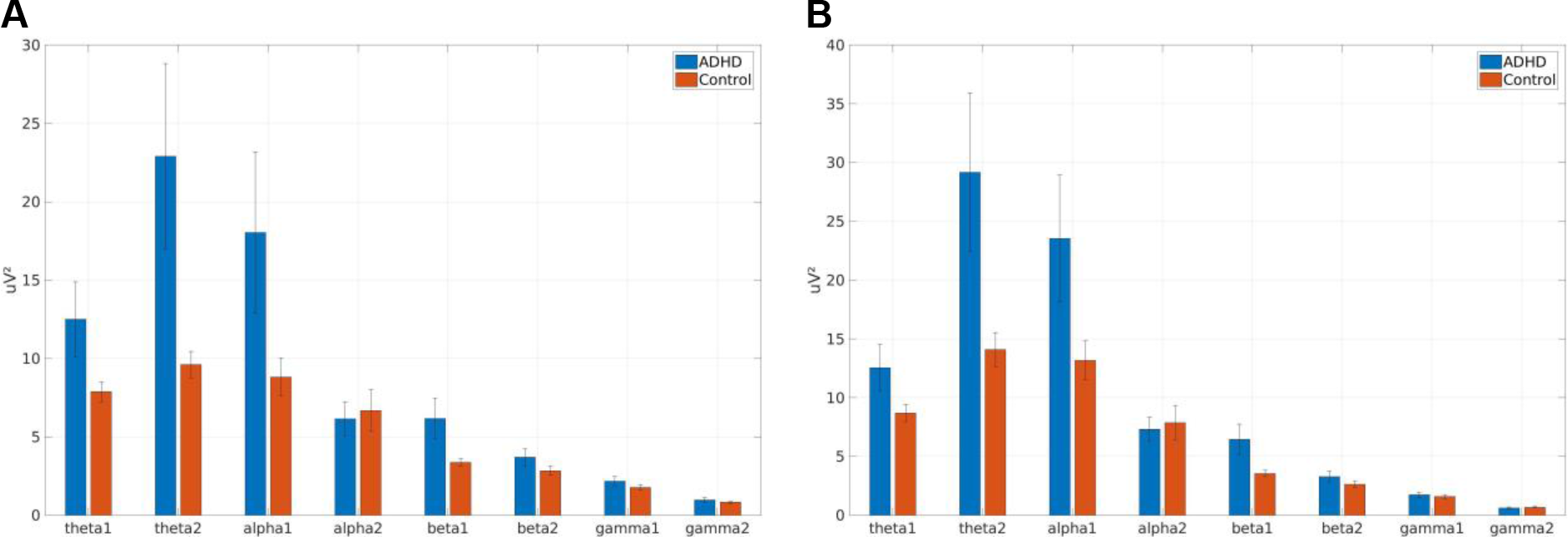
Power grand averages for ADHD and control population in A) Eyes open and B) eyes closed conditions.

Figure 6 presents the grand average and standard error of the mean of the EC-EO power ratio metric defined in 3.4. In both populations we observe a power increase in EC in low frequencies (theta and alpha), while in high frequencies (beta and gamma) the power increases in EO. Statistical differences between populations, as seen in table 2D, were not found at any band or electrode.

**Fig. 5.**
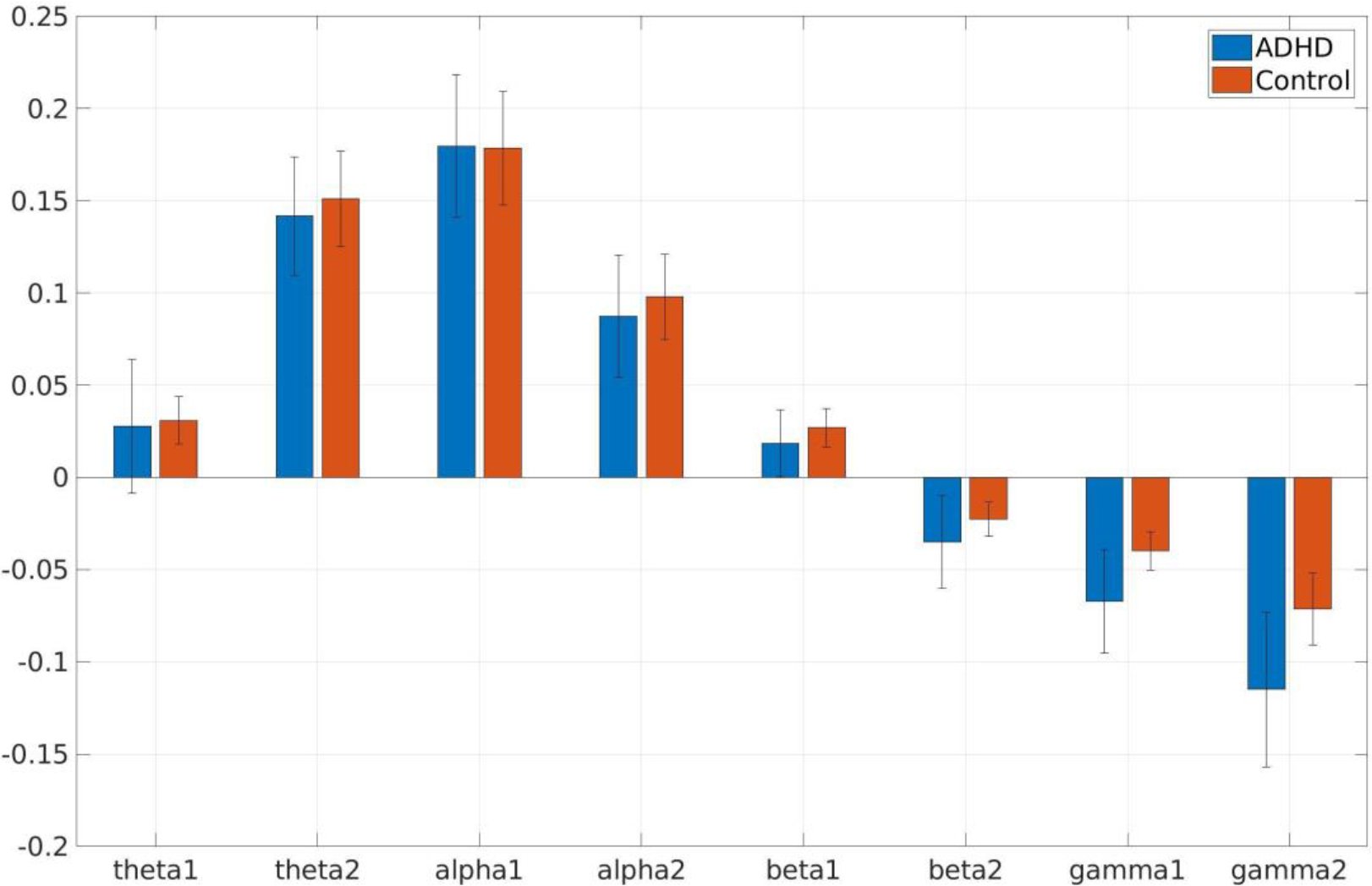
EC-EO power ratio grand averages in ADHD and control populations.

## 5. DISCUSSION

The diagnostic of ADHD is currently established based on subjective behavioral measures. ADHD diagnosis is therefore biased by cultural, practice and experience factors of clinicians. In the last decades, it has been shown that children suffering from ADHD may have different neural organization, especially in central and frontal areas compared to their age-matched control population. This fact advocates for the use of non-invasive brain monitoring techniques, such as EEG, to quantify abnormal brain activity patterns. Despite great advances in the field of ADHD biomarker discovery, the current state-of-the-art methodologies, such as the well accepted stationary theta increase, beta decrease in ADHDs, have proved insufficiently accurate in many scenarios. Moreover, the non-stationary and chaotic nature of brain activity supports the use of non-stationary techniques capable of detecting abnormal signal dynamics.

ADHD, among other behavioral symptoms, is characterized by a child’s inability to sustain attention as well as inappropriate arousal levels. In this study, we propose to study brain activity disparity between low arousal/attention levels (resting eyes closed) and normal attention/arousal levels (resting eyes open) as introduced in section 1. Our hypothesis relies in the fact that the disparity between these two conditions may be altered in the ADHD population. To measure it we propose a machine-learning based methodology capable of learning dynamical differences between EC-EO conditions. An echo state network input is fed using the filtered temporal series of pairs of electrodes and trained to discriminate between these two. As the amplitude information of these series is removed through standardization, the neural network is believed only to capture the signal dynamics. In previous works we have proved that ESNs are capable of detecting generalized synchronization between two temporal time-series [55]. As we are in a similar scenario we expect ESN to, among other dynamics, detect synchronization variations among channels between EC and EO regimes. The difference between EO and EC has also been studied through a stationary approach that computes the power ratio between EC and EO.

We have analyzed EEG recordings of 21 ADHD children and 30 age-matched controls and studied statistical differences between these two groups. To this effect we have evaluated the performance of stationary power during resting EO and EC, stationary EC-EO power differences and the outcome of the CDSM and DCI features proposed here.

Stationary spectral analysis showed an EEG slowing both for EC and EO in the ADHD population, where grand average alpha peak frequency was shifted towards slower frequencies in the ADHD population (Figure 4). A generalized slowing in brain activity can be linked to neurodegenerative and neurodevelopmental disorders and has been largely reported in the ADHD population [58, 59]. Stationary power delivers a statistically significant increase of theta in the ADHD population both in EC and EO conditions (Table 2). ADHD children have been largely reported to a show fairly consistent fronto-central theta increase in resting state conditions that has traditionally been associated to a hypo-arousal state [60, 61]. The outcome of this spectral stationary analysis confirm the reliability of the data-set, as two of most reported findings in the literature could be replicated.

The EO/EC band-power stationary analysis did not deliver statistical differences between the ADHD and control population. On the other hand, when analyzing the dynamical variations measured through the hereby proposed Dynamical Connectivity Index, statistically significant differences were found in the theta and beta bands. Abnormalities in these bands have been largely reported in the ADHD population as previously stated in section 1. This study confirms that not only the stationary activity of the brain, but also its temporal dynamics, may be affected at these bands. We have demonstrated that our trained network discriminates better between EC and EO regimes in the ADHD population. This finding indicates an abnormal disparity between low and normal attention/arousal conditions in ADHDs. Changes between conditions were only statistically significant for the DCI feature, fact that advocates for the employment of nonlinear analysis such as ESN, capable of discriminating between complex patterns in EEG time-series.

Compared to stationary power analysis, the DCI metric proposed here provides larger statistical significance between groups, as seen in Table 2. While DCI provides 11 electrode-band combinations with null hypothesis probability values below 1%, the EO and EC stationary power together provides 8. Whereas these differences were found in theta2 in the stationary power analysis both for EC and EO, DCI provided statistically significant differences below 1% in theta1 and beta1.

Reservoir-computing methodologies, and in particular ESNs, have proved to be an effective technique for EEG feature extraction [39]. Although ESNs have been applied to other EEG analysis scenarios including author’s work in Parkinson’s disease prognosis [37], this is the first time, to our best knowledge, that it has been used in the ADHD field. The proposed methodology is however not only tied to ADHD characterization. The approach of measuring the training error of a recurrent neural network trained to classify sequences of physiological states concatenated within a continuous time-series can be applied to other biomarker discovery modalities. Authors plan in the future to apply this procedure in the field of autism disorder characterization.

## 6. ACKNOWLEDGMENTS

We specially acknowledge the participants in this study. Funding was provided by the European Commission through the STIPED project under the Horizon 2020 Program (H2020, Grant agreement number 731827). JGO was financially supported by the Spanish Ministry of Economy and Competitiveness and FEDER (project FIS2015-66503-C3-1-P and Maria de Maeztu programme, MDM-2014-0370), the Generalitat de Catalunya (project 2017 SGR 1054), and the ICREA Academia program. We thank Patricia Teruel and Marta García Pérez for their assistance in the EEG recordings.

## REFERENCES

1. American Psychiatric Association (1994) Diagnostic and statistical manual of mental disorders, 4th ed. American Psychiatric Association, Washington, DC

2. Cormier, E. (2008). Attention deficit/hyperactivity disorder: a review and update. Journal of pediatric nursing, 23(5), 345–357.

3. American Psychiatric Association. (2000). Diagnostic criteria from dsM-iV-tr. American Psychiatric Pub.

4. Rowland, A. S., Lesesne, C. A., & Abramowitz, A. J. (2002). The epidemiology of attention-deficit/hyperactivity disorder (ADHD): a public health view. Mental retardation and developmental disabilities research reviews, 8(3), 162–170.

5. American Psychiatric Association, 2013

6. Willcutt, E. G. (2012). The prevalence of DSM-IV attention-deficit/hyperactivity disorder: a meta-analytic review. Neurotherapeutics, 9(3), 490–

7. Ravens-Sieberer, U., Wille, N., (2008b) Prevalence of mental health problems among children and adolescents in Germany: results of the BELLA study within the National Health Interview and Examination Survey. European child & adolescent psychiatry, 17 Suppl 1, 22–3

8. American Psychiatric Association. (2013). Diagnostic and statistical manual of mental disorders (5th ed.). Washington, DC: American Psychiatric Association.

9. Polanczyk, G. V., Willcutt, E. G., Salum, G. A., Kieling, C., & Rohde, L. A. (2014). ADHD prevalence estimates across three decades: an updated systematic review and meta-regression analysis. International Journal of Epidemiology, 43(2), 434–442. doi:10.1093/ije/dyt261

10. Crippa, A. et al. (2017). The Utility of a Computerized Algorithm Based on a Multi-Domain Profile of Measures for the Diagnosis of Attention Deficit/Hyperactivity Disorder. Frontiers in Psychiatry, 8 (189). doi:10.3389/fpsyt.2017.00189

11. Buyck, I., & Wiersema, J. R. (2014). Resting electroencephalogram in attention deficit hyperactivity disorder: developmental course and diagnostic value. Psychiatry research, 216(3), 391–397

12. Snyder SM, Rugino, TA, Hornig, M, Stein, MA. Integration of an EEG biomarker with a clinician’s ADHD evaluation. Brain Behav. 2015;5: 1–17. doi:10.1002/brb3.330

13. Geir, Ogrim, Juri Kropotov KH. The QEEG theta/beta ratio in ADHD and normal controls: Sensitivity, specificity and behavioral correlates. Psychiatry Res. 2012;198: 482–488. doi:10.1016/j.psychres.2011.12.041

14. Buyck, I, Wiersema, JR. Resting electroencephalogram in attention deficit hyperactivity disorder: Developmental course and diagnostic value. Psychiatry Res. 2014;216: 391–397. doi:10.1016/j.psychres.2013.12.055

15. Clarke, AR, Barry, RJ, McCarthy, R, Selikowitz, M, Johnstone, SJ, Hsu, CI, et al. Coherence in children with Attention-Deficit/Hyperactivity Disorder and excess beta activity in their EEG. Clin Neurophysiol. 2007;118: 1472–1479. doi:10.1016/j.clinph.2007.04.006

16. Theta Beta Ratio Ref

17. Gloss, D., Varma, J. K., Pringsheim, T., & Nuwer, M. R. (2016). Practice advisory: The utility of EEG theta/beta power ratio in ADHD diagnosis Report of the Guideline Development, Dissemination, and Implementation Subcommittee of the American Academy of Neurology. Neurology, 87(22), 2375–2379.

18. Snyder, S. M., Rugino, T. A., Hornig, M., & Stein, M. A. (2015). Integration of an EEG biomarker with a clinician’s ADHD evaluation. Brain and behavior, 5(4).

19. Ogrim, G., Kropotov, J., & Hestad, K. (2012). The quantitative EEG theta/beta ratio in attention deficit/hyperactivity disorder and normal controls: sensitivity, specificity, and behavioral correlates. Psychiatry research, 198(3), 482–488.

20. Liechti, M. D., Valko, L., Müller, U. C., Döhnert, M., Drechsler, R., Steinhausen, H. C., & Brandeis, D. (2013). Diagnostic value of resting electroencephalogram in attention-deficit/hyperactivity disorder across the lifespan. Brain topography, 26(1), 135–151.

21. Fabien, Lotte. A Tutorial on EEG Signal Processing Techniques for Mental State Recognition in Brain-Computer Interfaces. Eduardo Reck Miranda; Julien Castet. Guide to Brain-Computer Music Interfacing, Springer, 2014.

22. Coyle, D. (2009). Neural network based auto association and time-series prediction for biosignal processing in brain-computer interfaces. Computational Intelligence Magazine, IEEE, 4(4), 47–59.

23. Lotte, F, Congedo, M, Lecuyer, A, Lamarche, F, Arnaldi, B (2007) A review of classification algorithms for EEG-based brain-computer interfaces. Journal of Neural Engineering 4:R1–R13

24. Lindsley, D. B. (1988). Activation, arousal, alertness, and attention. In States of brain and mind (pp. 1–3). Birkhäuser Boston.

25. Barbaro, K., Clackson, K., & Wass, S. V. (2017). Infant attention is dynamically modulated with changing arousal levels. Child development, 88(2), 629–639.

26. Sonuga-Barke EJ, Wiersema, JR, van der Meere JJ, Roeyers, H (2010) Context-dependent dynamic processes in attention deficit/hyperactivity disorder: differentiating common and unique effects of state regulation deficits and delay aversion. Neuropsychol Rev 20(1):86–102.

27. Loo, S. K., Hale, T. S., Macion, J., Hanada, G., McGough, J. J., McCracken, J. T., & Smalley, S. L. (2009). Cortical activity patterns in ADHD during arousal, activation and sustained attention. Neuropsychologia, 47(10), 2114–2119.

28. Geissler, J., Romanos, M., Hegerl, U., & Hensch, T. (2014). Hyperactivity and sensation seeking as autoregulatory attemps to stabilize brain arousal in ADHD and mania? Attention Deficit Hyperactivity Disorder, 6 (159–173). doi:10.1007/s12402-014-0144-z

29. Halperin, JM, Schulz, KP (2006) Revisiting the role of the prefrontal cortex in the pathophysiology of attention-deficit/hyperactivity disorder. Psychol Bull 132(4):560–581

30. Barry, R. J., & De Blasio, F. M., (2017). EEG Differences between Eyes-Closed and Eyes-Open Resting Remain in Healthy Ageing. Biological Psychology.

31. Ramachandran, V. S. (2002). Encyclopedia of the Human Brain, Four-Volume Set. Academic Press.

32. Aminoff, M. J. (2012). Aminoff’s Electrodiagnosis in Clinical Neurology E-Book. Elsevier Health Sciences.

33. Barry, R. J., Clarke, A. R., Johnstone, S. J., Magee, C. A., & Rushby, J. A. (2007). EEG differences between eyes-closed and eyes-open resting conditions. Clinical Neurophysiology, 118(12), 2765–2773.

34. Coone, A., & Schrauwen, B. (2011). Single trial classification of the P300 evoked potential using Reservoir Computing. In 1st Joint WIC/IEEE SP Symposium on Information Theory and Signal Processing in the Benelux.

35. Gouy-Pailler, C., Sebag, M., Souloumiac, A., & Larue, A. (2010, August). Ensemble learning for brain computer-interface using uncooperative democratic echo state communities. In Cinquième conférence plénière française de Neurosciences Computationnelles,” Neurocomp'10”.

36. Buteneers, P., Verstraeten, D., van Mierlo, P., Wyckhuys, T., Stroobandt, D., Raedt, R., … & Schrauwen, B. (2011). Automatic detection of epileptic seizures on the intra-cranial electroencephalogram of rats using reservoir computing. Artificial intelligence in medicine, 53(3), 215–223.

37. Ruffini, G., Ibañez, D., Castellano, M., Dunne, S., & Soria-Frisch, A. (2016, September). EEG-driven RNN classification for prognosis of neurodegeneration in at-risk patients. In International Conference on Artificial Neural Networks (pp. 306–313). Springer, Cham.

38. Ayyagari, S. S., Jones, R. D., & Weddell, S. J. (2014, August). EEG-based event detection using optimized echo state networks with leaky integrator neurons. In 2014 36th Annual International Conference of the IEEE Engineering in Medicine and Biology Society (pp. 5856–5859). IEEE

39. D. Ibanez-Soria, A. Soria-Frisch, J. Garcia-Ojalvo, G. Ruffini (2018). Echo State Networks Ensemble for SSVEP Dynamical Online Detection. bioArxiv.

40. Bishop, C. M. (1995). Neural networks for pattern recognition. Oxford university press.

41. Duda, R. O., Hart, P. E., & Stork, D. G. (2012). Pattern classification. John Wiley & Sons.

42. Yegnanarayana, B. (2009). Artificial neural networks. PHI Learning Pvt. Ltd.

43. Schrauwen, B., Verstraeten, D., & Van Campenhout, J. (2007). An overview of reservoir computing: theory, applications and implementations. In Proceedings of the 15th European Symposium on Artificial Neural Networks. p. 471-482 2007 (pp. 471–482).

44. Jaeger, H.: The “echo state” approach to analysing and training recurrent neural networks. Tech. Rep. GMD Report 148, German National Research Center for Information Technology (2001).

45. Natschläger, T., Maass, W., & Markram, H. (2002). The” liquid computer": A novel strategy for real-time computing on time series. Special issue on Foundations of Information Processing of TELEMATIK, 8(LNMC-ARTICLE-2002-005), 39–43.

46. Lukoševičius, M. (2012).A practical guide to applying echo state networks. In Neural networks: Tricks of the trade (pp. 659–686). Springer Berlin Heidelberg

47. Yildiz, I. B., Jaeger, H., & Kiebel, S. J. (2012). Re-visiting the echo state property. Neural networks, 35, 1–9

48. David Verstraeten, Joni Dambre, Xavier Dutoit, and Benjamin Schrauwen. Memory versus non-linearity in reservoirs. In Proc. Int Neural Networks (IJCNN) Joint Conf, pages 1–8, 2010.

49. Fabian Triefenbach, Azarakhsh Jalalvand, Benjamin Schrauwen, and Jean-Pierre Martens. Phoneme recognition with large hierarchical reservoirs. In Advances in Neural Information Processing Systems 23 (NIPS 2010), pages 2307–2315. MIT Press, Cambridge, MA, 201

50. Wyffels, F., Schrauwen, B., & Stroobandt, D. (2008, September). Stable output feedback in reservoir computing using ridge regression. In International conference on artificial neural networks (pp. 808–817). Springer, Berlin, Heidelberg.

51. Arns, M., Conners, C. K., & Kraemer, H. C. (2012). A decade of EEG theta/beta ratio research in ADHD: a meta-analysis. Journal of attention disorders, 1087054712460087.

52. Jurcak, V., Tsuzuki, D., & Dan, I. (2007). 10/20, 10/10, and 10/5 systems revisited: their validity as relative head-surface-based positioning systems. Neuroimage, 34(4), 1600–1611.

53. Teplan, M. (2002). Fundamentals of EEG measurement. Measurement science review, 2(2), 1–11.

54. Yegnanarayana, B. (2009). Artificial neural networks. PHI Learning Pvt. Ltd.

55. Ibanez-Soria, D., Garcia-Ojalvo, J., Soria-Frisch, A., & Ruffini, G. (2017). Detection of Generalized Synchronization using Echo State Networks. arXiv preprint arXiv:1710.08286.

56. Bohm, G., & Zech, G. (2010). Introduction to statistics and data analysis for physicists. DESY.

57. Wilcoxon, F. (1945). Individual comparisons by ranking methods. Biometrics bulletin, 1(6), 80–83.

58. Vollebregt, M. A., van Dongen-Boomsma, M., Slaats-Willemse, D., Buitelaar, J. K., & Oostenveld, R. (2015). How the individual alpha peak frequency helps unravel the neurophysiologic underpinnings of behavioral functioning in children with attention-deficit/hyperactivity disorder. Clinical EEG and neuroscience, 46(4), 285–291.

59. Lansbergen, M. M., Arns, M., van Dongen-Boomsma, M., Spronk, D., & Buitelaar, J. K. (2011). The increase in theta/beta ratio on resting-state EEG in boys with attention-deficit/hyperactivity disorder is mediated by slow alpha peak frequency. Progress in Neuro-Psychopharmacology and Biological Psychiatry, 35(1), 47–52

60. Snyder, SM, Hall, JR. A meta-analysis of quantitative EEG power associated with attention-deficit hyperactivity disorder. J Clin Neurophysiol. 2006;23:440–455.

61. Loo, S. K., & Makeig, S. (2012). Clinical utility of EEG in attention-deficit/hyperactivity disorder: a research update. Neurotherapeutics, 9(3), 569–587.

